# Diagnostic efficacy of hand-held digital refractometer for determining total serum protein in indigenous sheep of Pakistan

**DOI:** 10.1101/2023.11.15.567327

**Authors:** Madiha Sharif, Mushtaq Hussain Lashari, Umer Farooq, Musadiq Idris, Muhaammad Abrar Afzal

## Abstract

The study was designed to ascertain the diagnostic efficacy of hand-held digital refractometer in determining total protein. The Sipli sheep (n=141) were grouped as per gender (females=99, males=29) and age (G1=up till 1 year, G2=from 1 to 2 years, G3=above 2 years). The results regarding the overall mean (±SE) values and RIs for the TPs attained through serum chemistry analyzer (TP1) and hand-held digital refractometer (TP2) were non-significantly (P≤0.05) different (59.2±1.6g/L and 59.8±0.5g/L, respectively). However, the RIs were quite different between the two TPs being 45.1-95.7g/L and 57.0-67.0g/L for TP1 and TP2, respectively. Similar results were seen for gender-wise and group-wise results. On the contrary, the results regarding correlation coefficient and logilinear regression showed a negative correlation between the two TPs (r=-0.0244) with an adjusted r-square of 0.059 (5.9% probability). Furthermore, the results for Cronbach alpha and intraclass correlation coefficient between TP1 and TP2 showed that the values for single measure and average values were lower between TP1 and TP2 being - 0.135 and −0.313. Bland and Altman test between TP1 and TP2 also showed a weak level of agreement between the two methods of detecting TP. A proportional bias on the distribution of data around the mean difference line was noticed between TP1 and TP2 (Mean= 0.5; 95% CI= 39.8 to −40.9) with a standard deviation of biasness being 20.58. In a nutshell, the hand-held digital refractometer cannot be used as an on-farm POCT device for determining serum TP in sheep. However, certain other models of refractometers with higher sensitivity and specificity may be utilized in future studies to establish these conclusions for other species of livestock.

## Introduction

The diagnostic/prognostic tests performed out of the laboratory and near/at the site of patient are termed as point-of-care-tests (POCTs) as per the definition provided by the International Standard ISO 22870 [1]. These tests have a rapid turnaround time, and assist in rapid decision-making and faster care of the patient. In human medical practices they are dubbed as bed-side tests, near-patient testing, and patient-focused testing, whereas in veterinary medical diagnostics, they are termed as cow-side tests, on-farm tests, barn-side tests or flock-side tests [2,3]. On human medical side, there are various POCTs presently in vogue for detecting glucose, hemoglobin, blood gases, electrolytes, cardiac enzymes and drug metabolites. In order to attain reliable results of surveillance, outbreak or analytic testing from the POCTs, it is vital that they are kept under strict regulatory oversight, quality control, and periodic comparison/validation through gold-standard techniques. Regarding veterinary medical diagnostics, there is still a paucity of devising and validating POCT devices for field use. It has been estimated that the market of veterinary POCTs was around 2.15 billion dollars with a prediction of 12.3% growth during 2021 to 2030 [4]. Evolving technologies and increased market needs are the key reasons behind this escalatory pattern of POCTs in order to efficiently respond the new and emerging diseases.

Since their introduction in 1960s, the hand-held digital refractometers are being used extensively as POCT devices for determining Brix%, salinity, specific gravity, and total proteins in various bodily fluids. These instruments measure the angle of refraction between air and an aqueous solution, providing rapid and inexpensive determination of solutes in the fluids. These devices have widely been used in veterinary medical diagnostics for assessing failure in passive transfer (FPT) through determining IgGs in sows [5], dogs [6] and cattle calves [7, 8]. Similarly, they have also been used for determining serum total proteins (TP) in various species with varying sensitivity and specificity as compared to other gold-standard techniques [9-11]. However, to the best of knowledge, the diagnostic efficacy of hand-held digital refractometers for determining TP in indigenous sheep of Pakistan has not yet been unearthed.

There are about 17 indigenous sheep breeds being reared in Pakistan in different geo-ecological zones. It has been claimed that these breeds probably originated from urial (*Ovisvignei*), the wild sheep of Afghanistan, Baluchistan and Central Asia [12]. Sipli breed of sheep is a thin-tailed indigenous sheep breed of Pakistan with a relatively long tail. Small number of this breed (n=260) is being maintained at two institutional farms in South Punjab, Pakistan. It is a medium-sized sheep breed with an average body weight of 32.8kg for males and 29.2kg for females, and has a daily milk yield of 0.2-0.4 L [12,13]. It has white body coat with white or light brown head/ears. Its head is medium sized and has a flat nose with ears reaching about 15 cm long [14]. It is mostly reared for mutton and wool purposes by the nomadic herders of Bahawalpur, Bahawalnagar and Rahim-Yar-Khan-the three cities which lay in the middle of the Cholistan desert, (Southern Punjab) Pakistan. The present research work is the first of its kind being reported for this sheep breed with an objective to ascertain the diagnostic efficacy of hand-held digital refractometer in determining TP as compared to serum chemistry analyzer.

## Materials and methods

### Geo-location of study

The present study was carried out simultaneously at the Livestock Farm, Faculty of Veterinary and Animal Sciences (FV&AS), The Islamia University of Bahawalpur (IUB), Pakistan and Post-graduate Lab of Physiology, IUB. The climate of Cholistan desert is arid and semi-arid tropical; the average temperature of Cholistan desert is 28.33°C, average rainfall of Cholistan desert is up to 180mm [15].

### Experimental animals

The Sipli breed of sheep (n=141) are being reared at the Livestock Farm of FV&AS, IUB, Pakistan under intensive farming system were incorporated in the present study. The animals under study were grouped as per gender (females=99, males=29) and age (G1=up till 1 year, G2=from 1 to 2 years, G3=above 2 years). The animals are sent for grazing early morning. In the evening the feeding of animals includes fresh-cut and chopped seasonal fodder along with concentrate ration containing about 15% crude protein. In addition, maize silage and wheat straw is offered depending on need as and when required. The fresh clean drinking water remains available all the time. The animals have been assigned tag numbers in order to collect data.

### Blood collection and analyses

Approximately 5mL blood sample were collected from each experimental animal. Bleeding was conducted once with a total of 141 blood samples. The blood was collected aseptically from the jugular vein using a 5mL disposable syringe in yellow-capped vacutainers containing silica and a polymer gel for serum separation. The vacutainers were centrifuged at 3000rpm for 15 minutes by centrifuge machine (Centrifuge 800, China) for serum extraction. Serum was extracted in eppendorf tubes which were transported in ice-packs to the Post-graduate Lab Physiology, IUB for further analysis.

The serum TP from attained serum samples was analyzed by two methods:

a. **In-lab** the TP was determined through commercially available TP kit (Bioactive Diagnostic Systems, JTC, Cat No. 5-172, Germany) using semi-automated serum chemistry analyzer (Rayto-9600, China) as per instruction manual. The commercial kit had a sensitivity/limit of quantification 0.17g/dL (7.3g/L) and Linearity of up to 15g/dL (150g/L). The TP thus attained was termed as TP1 in this study.
b. **On-field** determination of TP was carried out using a hand-held digital refractometer (Serum Protein Tester, DR503, China) as per the instructions of the manufacturer. Briefly, after turning on the device, it was cleared with the distilled water and the sample plate was dried. A 0.2-0.3 mL sample was dripped on the plate, and the cover/lid was closed. Pressing the ‘read’ button one time provided the TP in g/dL. The devise had a performance range of 2.0-14.0g/dL, accuracy of ±0.2, and a resolution of 0.1. The TP thus attained was termed as TP2 in this study.

### Statistical analyses

Outliers in the data were inspected visually and upon finding 13 outliers, these were removed for further analyses. Hence, RIs were deduced for remaining data (n=128) keeping in view the guidelines provided by the American Society for Veterinary Clinical Pathology [16] using the Reference Value Advisor (freeware v.2.1: http://www.biostat.envt.fr/reference-value-advisor). Remaining tests were performed through Statistical Package for Social Sciences (Windows Version 12, SPSS Inc, Chicago, IL, USA). Mean (±SE) values for serum TP attained through serum chemistry analyzer (TP1) and through digital hand-held refractometer (TP2) were analyzed for difference through independent t-test. Pearson’s correlation coefficient and linear regression were implied to assess the level of relation between the two values. Three tests were implied to check the level of agreement between the two type of tests *viz*. Bland & Altman, Cronbach Alpha and Intraclass Coefficient.

## Results

The results regarding the overall mean (±SE) values and overall RIs for the TP1 (attained through serum chemistry analyzer) and TP2 (attained through hand-held digital refractometer) are given in Table 1. Both the mean (±SE) values (for TP1 and TP2) were non-significantly (P≤0.05) different with mean values of 59.2±1.6g/L and 59.8±0.5g/L, respectively. However, the RIs were quite different between the two TPs being 45.1-95.7g/L and 57.0-67.0g/L for TP1 and TP2, respectively. Similar results were seen for gender-wise and group-wise results being non-significantly (P≤0.05) different between all study groups, but with wide range of RIs. Group-wise results for TP1 and TP2 are given in Table 2 and 3, respectively.

**Table 1.** Overall mean (±SE), median, interquartile range, minimum, maximum, 25th to 95th percentile of reference interval (RI) and 95% confidence interval (CI) for serum total protein in Sipli sheep (n=128)

| Attribute | Mean (±SE) | Median (IQR) | Range (Min-Max) | RI (25<sup>th</sup> to 95<sup>th</sup>) | 95% CI |
| --- | --- | --- | --- | --- | --- |
| TP1 (g/L) | $59.2 \pm 1.6$ | 58.5(26.3) | 78.3(21.2-99.5) | 45.1-95.7 | 56.0-62.5 |
| TP2 (g/L) | $59.8 \pm 0.5$ | 60.0(6.7) | 47.0(34.0-81.0) | 57.0-67.0 | 58.8-60.7 |

TP1: Total protein estimated through serum chemistry analyzer; TP2: Total protein attained through hand-held digital refractometer

**Table 2.** Mean (±SE), median, interquartile range, minimum, maximum, 25th to 95th percentile of reference interval (RI) and 95% confidence interval (CI) for serum total protein (g/dL) attained through serum chemistry analyzer as affected by sex and age in Sipli sheep (n=128)

| Groups | Mean±SE | Median (IQR) | Range (Min-Max) | RI<br>(25<sup>th</sup> to 95<sup>th</sup>) | 95% CI |
| --- | --- | --- | --- | --- | --- |
| Females (n=99) | 59.3±1.8 | 58.2 | 78.3(21.2-99.5) | 44.7-95.8 | 55.6-63.1 |
| Males (n=29) | 58.7±3.2 | 58.9 | 70.6(25.0-95.7) | 45.7-93.1 | 52.1-65.4 |
| G1 (n=35) | 58.1±3.1 | 59.1 | 76.5(21.2-97.7) | 44.6-96.5 | 51.6-64.6 |
| G2 (n=63) | 58.9±2.4 | 57.8 | 77.2(22.3-99.5) | 43.1-95.7 | 54.0-63.8 |
| G3 (n=30) | 61.0±2.9 | 58.5 | 61.3(34.4-95.8) | 45.8-91.7 | 54.9-67.1 |
| Overall (n=128) | 59.2±1.6 | 58.5(26.3) | 78.3(21.2-99.5) | 45.1-95.7 | 56.0-62.5 |

**Table 3.** Mean (±SE), median, interquartile range, minimum, maximum, 25th to 95th percentile of reference interval (RI) and 95% confidence interval (CI) for serum total protein (g/dL) attained through hand-held digital refractometer as affected by sex and age in Sipli sheep (n=128)

| Groups | Mean±SE | Median (IQR) | Range (Min-Max) | RI<br>(25<sup>th</sup> to 95<sup>th</sup>) | 95% CI |
| --- | --- | --- | --- | --- | --- |
| Females (n=99) | 59.4±0.6 | 60.0 | 47.0(34.0-81.0) | 56.0-67.0 | 58.2-60.6 |
| Males (n=29) | 60.9±0.7 | 61.0 | 17.0(51.0-68.0) | 58.5-67.5 | 59.4-62.4 |
| G1 (n=35) | 59.8±1.3 | 61.0 | 47.0(34.0-81.0) | 57.0-72.2 | 57.1-62.5 |
| G2 (n=63) | 59.3±0.6 | 60.0 | 22.0(47.0-69.0) | 56.0-66.8 | 58.1-60.5 |
| G3 (n=30) | 60.6±0.6 | 60.0 | 17.0(51.0-68.0) | 58.9-67.4 | 59.2-62.0 |
| Overall (n=128) | 59.8±0.5 | 60.0(6.7) | 47.0(34.0-81.0) | 57.0-67.0 | 58.8-60.7 |

The results regarding correlation coefficient and logilinear regression (Fig 1) between TP1 and TP2 showed a negative correlation between the two attributes (r=-0.0244) and an adjusted r-square of 0.059 (5.9% probability).

**Fig 1.**
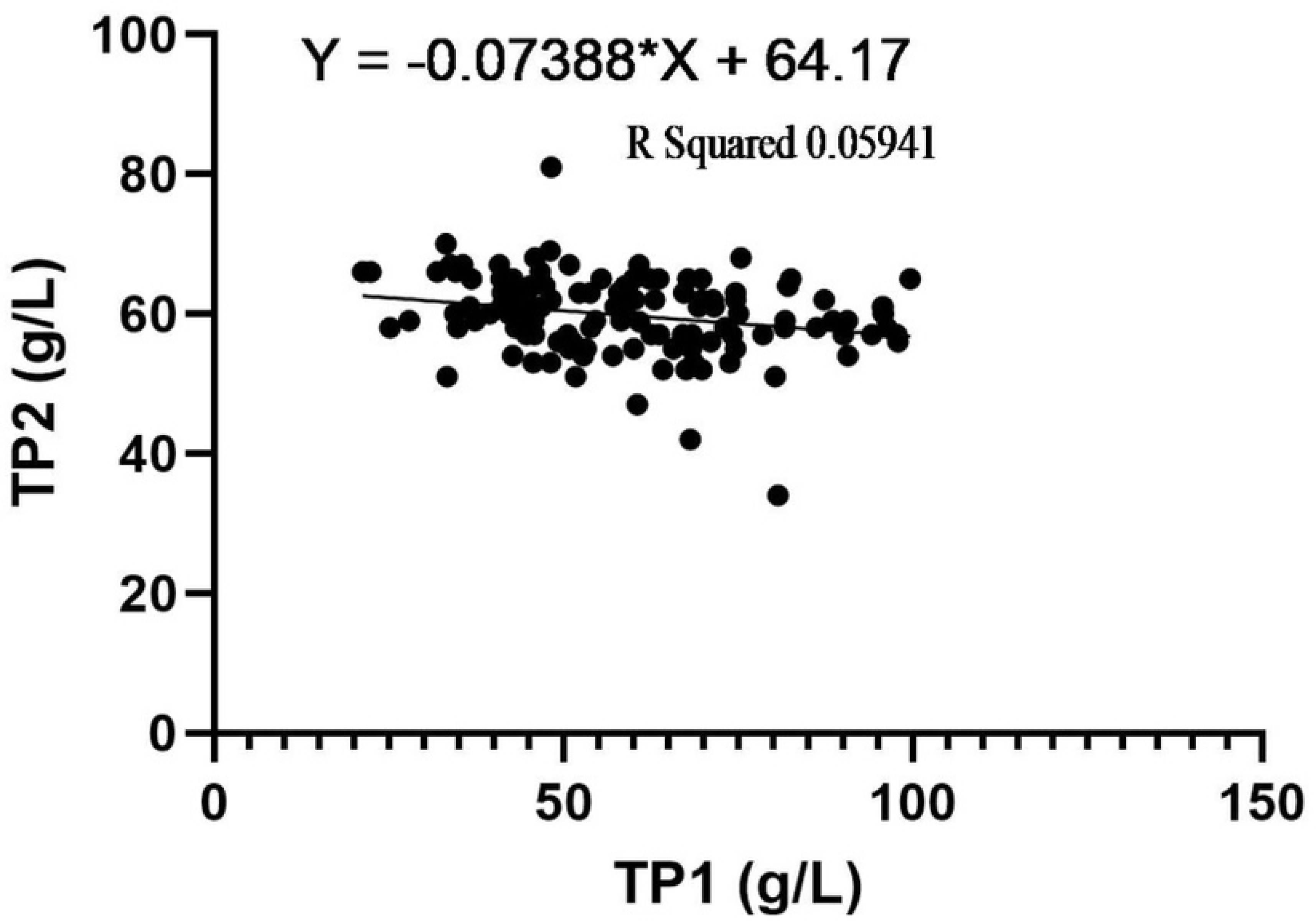
Scatterplot for logilinear regression between TP1 (attained through serum chemistry analyzer) and TP2 (attained through hand-held digital refractometer)

Results for Cronbach alpha and intraclass correlation coefficient between TP1 and TP2 are given in Table 4. Both the values for single measure and average values were lower between TP1 and TP2 being −0.135 and −0.313, respectively.

**Table 4.** Cronbach alpha and intraclass correlation between TP1 (attained through serum chemistry analyzer) and TP2 (attained through hand-held digital refractometer)

| TP1 vs TP2 |  |  |  |
| --- | --- | --- | --- |
| Intraclass Correlation |  | 95% CI | Cronbach Alpha |
| Single Measure | -0.135 | -0.301-0.038 | -0.313 |
| Average Measures | -0.313 | -0.862-0.074 |  |

TP1: Total protein estimated through serum chemistry analyzer; TP2: Total protein attained through hand-held digital refractometer

Similarly, Bland and Altman chart between TP1 and TP2 (Fig 2) showed a weak level of agreement. A proportional bias on the distribution of data around the mean difference line was noticed between TP1 and TP2 (Mean= 0.5; 95% CI= 39.8 to −40.9) with an S.D. of biasness being 20.58.

**Fig 2.**
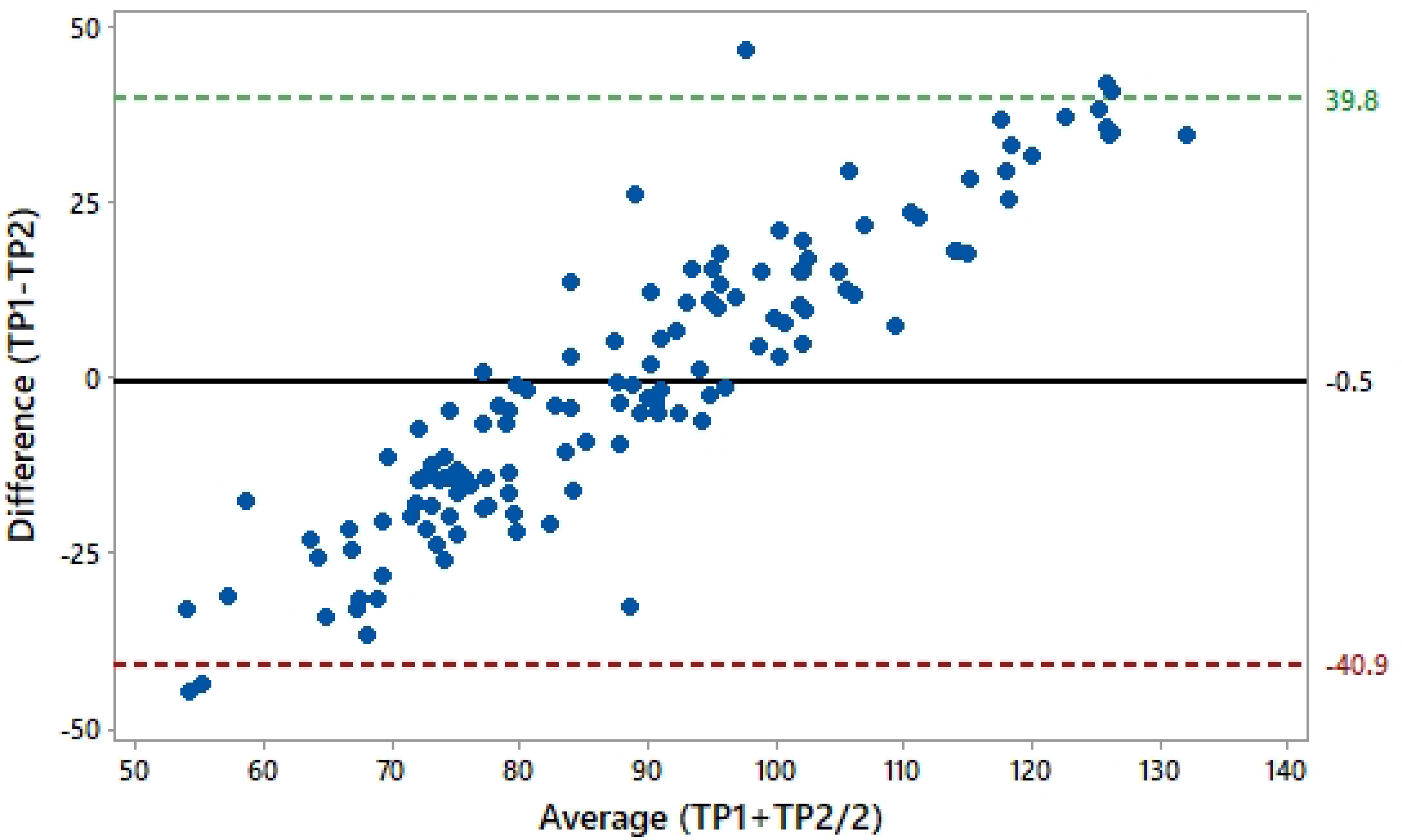
Scatterplot of Bland and Altman test between difference of total protein determined through serum chemistry analyzer and through hand-held digital refractometer (TP1-TP2) and average of both TPs (TP1+TP2/2) in Sipli sheep (n= 141). Black line indicates mean difference (−0.5) whereas the upper and lower dotted lines indicate upper (39.8) and lower (−40.9) values for 95% CI, respectively (SD of Bias 20.58)

## Discussion

The present work is the first of its kind being reported for indigenous Sipli breed of sheep from Pakistan which was conducted with an aim to assess diagnostic efficacy of hand-held digital refractometer for determining serum TP, as compared to the serum chemistry analyzer which is considered as the gold-standard technique in this study. Though globally acclaimed and in vogue, the digital hand-held refractometer failed to reveal satisfactory results for determining serum TP of the sheep in the present study as compared to those attained through serum chemistry analyzer.

In the present study, the mean values of 59.2±1.6g/L and 59.8±0.5g/L attained through serum chemistry analyzer using commercial kit, and through hand-held digital refractometer are in line with those reported for three indigenous sheep of Iraq being 62.0g/L [17]. Similar values of 58.2g/L have been reported for White Dorper and Suffolk sheep using Biuret method of TP determination [18]. Another study has also reported similar value of 6.0g/L using Biuret method for Dorper sheep of Brazil [19]. Higher values of 7.2g/L have been reported for free-ranging desert big-horn sheep [20] and Merino sheep [21] using automated serum chemistry analyzer. Variation in breed and method of TP determination may be attributed to these higher values of TP.

Regarding the RIs of TP of sheep serum in the present study, attained through two methods *i*.*e*. serum chemistry analyzer and hand-held digital refractometer, it was noticed that the RIs were quite different between the two TPs being 45.1-95.7g/L and 57.0-67.0g/L for TP1 and TP2, respectively. Comparing these results with prior studies, it was revealed that the values attained through hand-held refractometer were above and far beyond those reported for sheep using either the automated chemistry analyzers or Biuret method of TP determination, whereas the values attained in the present study through serum chemistry analyzer were in line with prior research reports [21-23]. The difference in these results indicates poor diagnostic efficacy of hand-held digital refractometer for determining TP in sheep serum.

Literature is rife with studies in which the digital refractometers (especially Brix refractometers) have successfully been used for determining serum immunoglobulins (IGs) in various species such as canines [6], bovines [7, 24, 25], porcine [26], ovine [27], and equine [28]. All these prior studies have validated hand-held digital refractometers for determining IGs in serum and have reported satisfactory sensitivity, specificity, negative predictive value and positive predictive values for this device dubbin it a reliable on-farm, cow-side POCT in contrary to our results.

Apart from correlation and regression which gave a weak association between the two methods of TP determination (r=-0.0244, adjusted r-square=0.059/5.9% probability), the present study utilized three tests (Bland & Altman, Cronbach Alpha and Intraclass Coefficient) for ascertaining level of agreement between the two methods using for determining TP *i*.*e*. through serum chemistry analyzer and hand-held digital refractometer. All these tests indicated a poor level of agreement between the two studied tests. The Bland & Altman test is considered as a gold-standard test for checking level of agreement between two different techniques measuring one similar variable [29-32]. For the present study, this test revealed a proportional bias on the distribution of data around the mean difference line between TP1 and TP2 (Mean= 0.5; 95% CI= 39.8 to −40.9) with an S.D. of biasness being 20.58.

Considering the results of the above study, it seems inevitable to conclude that hand-held digital refractometer cannot be used as an on-farm POCT device for determining serum TP in sheep. However, certain other models of refractometers with higher sensitivity and specificity may be utilized in future studies to establish these conclusions.

## Author contributions

**Conceptualization**: Umer Farooq and Mushtaq Hussain Lashari

**Data curation:** Madiha Sharif

**Formal analysis:** Musadiq Idris

**Methodology and software:** Muhammad Abrar Afzal

**Writing:** Umer Farooq

**Review and editing:** Umer Farooq and Madiha Sharif

